# Chikungunya virus infection impairs the function of osteogenic cells

**DOI:** 10.1101/2020.04.15.044065

**Authors:** Enakshi Roy, Wen Shi, Bin Duan, St Patrick Reid

**Affiliations:** Department of Pathology & Microbiology, University of Nebraska Medical Center, Omaha, NE 68198-5900, USA; Mary & Dick Holland Regenerative Medicine Program, Division of Cardiology, Department of Internal Medicine, University of Nebraska Medical Center, Omaha, NE 68198-5900, USA

**Keywords:** Chikungunya virus, mesenchymal stem cells, osteogenic differentiation

## Abstract

Chikungunya virus (CHIKV) is a positive-sense, single-stranded RNA virus, spread by the *Aedes* species (sp.) mosquitoes. Chikungunya virus (CHIKV) causes a condition characterized by high fever, headache, rash, and joint pain. Recent investigations reveal presence of bone lesions and erosive arthritis in the joints of CHIKV infected patients, indicating an association of bone pathology with CHIKV infection. However, the molecular mechanism underlying CHIKV-induced bone pathology remains poorly defined. Bone marrow derived mesenchymal stem cells (BMSCs) contribute to bone homeostasis by differentiating into osteogenic cells which later mature to form the bone. Disruption of osteogenic differentiation and function of BMSCs lead to bone pathologies. Studies show that virus infections can alter the properties and function of BMSCs. However, to date, pathogenesis of CHIKV infection in this context has not been studied. In the current study, we investigated the susceptibility of BMSCs and osteogenic cells to CHIKV and studied the effect of infection on these cells. To our knowledge, for the first time we report that CHIKV can productively infect BMSCs and osteogenic cells. We also observed a decreased gene expression of the major regulator of osteogenic differentiation, RUNX2 in CHIKV infected osteogenic cells. Furthermore, impaired functional properties of osteogenic cells i.e. decreased production and activity of alkaline phosphatase (ALP) and matrix mineralization were observed in the presence of CHIKV infection. Thus, we conclude that CHIKV likely impairs osteogenic differentiation of BMSCs indicating a possible role of BMSCs in altering bone homeostasis during CHIKV infection.

**Importance:** Presently, no vaccines or treatment options are available for CHIKV infection. Joint pain is one of the major concerns. Although studies have shown an association between bone pathology and infection, the molecular pathogenesis in context of bone pathology is poorly defined. Here, we demonstrate for the first time that BMSCs and BMSC-derived osteogenic cells are susceptible to CHIKV infection and infection likely alters function of the osteogenic cells. This study highlights altered osteogenic differentiation as a possible mechanism for causing the bone pathology observed in CHIKV pathogenesis.

## Introduction

Chikungunya virus (CHIKV) is a positive-sense, single-stranded RNA virus belonging to the *Togaviridae* family and *Alphavirus* genus [21]. Since mid-1900s, there have been outbreaks of CHIKV infection in Africa, Asia, and the Indian and Pacific Ocean region, with few reported cases within Europe [29]. Beginning in 2013, two independent CHIKV strains have been introduced in the Americas, in part due to travel from affected regions [27, 29]. CHIKV is transmitted by *Aedes* species (sp). mosquitoes and has infected millions of people annually causing CHIKV fever (CHKF) in the tropical and subtropical regions of the world [29]. CHIKF is characterized by a self-limiting acute stage, with symptoms of fever, headache, rash and arthralgia which lasts for 1-2 weeks [8]. In 30-40% of cases, there is a chronic stage where patients develop an incapacitating arthritis which may persist for months to years, and thereby imposes a burden on the population in terms of disability adjusted life years (DALY) [8, 11].

Recent studies identified bone lesions in the joints of CHIKV infected mice, indicating that CHIKV can cause bone pathologies [12]. In another study, mice infected with a similar arthritogenic alphavirus, Ross River virus (RRV) resulted in significant bone loss [7]. In humans, magnetic resonance imaging (MRI) studies revealed that CHIKV infection is associated with erosive arthritis [19]. Taken together these studies suggest alphavirus infection can affect bone homeostasis and thus contribute to arthritic-like conditions.

Mesenchymal stem cells (MSCs) are multipotent, non-hematopoietic stromal cells which can self-renew and differentiate into various cell lineages [28]. MSCs can be derived from umbilical cord blood, adipose tissue and bone marrow [28]. Bone marrow-derived MSCs (BMSCs) have trilineage differentiation potential and they can differentiate into osteogenic, chondrogenic or adipogenic cell lineage [18]. Osteogenic differentiation of BMSCs is important for bone homeostasis, and the inability of BMSCs to differentiate into the osteogenic cell lineage may lead to an imbalance in bone homeostasis and often causing bone pathology [26]. A few studies have shown that virus infection of BMSCs can affect the properties and function of these cells [22, 25].

In this study, we investigated the susceptibility of BMSCs and BMSC-derived osteogenic cells to CHIKV infection and the response of these cells to CHIKV infection. We hypothesized that CHIKV can infect BMSCs and affect the osteogenic differentiation of BMSCs. Our results show that CHIKV can productively infect BMSCs and BMSC-derived osteogenic cells. Interestingly, we observed a significant decrease in gene expression of the transcription factor and the major regulator of early osteogenic differentiation, RUNX2 in the presence of CHIKV infection [5]. More importantly, we observed that viral infection significantly impaired the function of the osteogenic cells as evidenced by the decrease in production and activity of alkaline phosphatase (ALP) and matrix mineralization i.e. production of calcium phosphate in the virus infected cells compared to mock-infected control [13]. Together, these findings indicate CHIKV can infect BMSCs and disrupt BMSC-derived osteogenic cell function.

## Results

### BMSCs are permissive to CHIKV infection

CHIKV infection has been associated with bone pathology, implying its role in disruption of bone homeostasis [12, 19]. BMSC-derived osteogenic differentiation is essential for bone homeostasis [26]. Recent studies show that viral infection can affect the function of BMSCs and BMSC-derived osteogenic cells [22, 25]. However, to date, it is unknown whether alphaviruses can infect BMSCs and disrupt osteogenic cell function. Permissivity of BMSCs to CHIKV infection was determined by infecting cells at multiplicity of infection (MOI) 1 (Fig.1A). To detect the presence of infection in BMSCs, immunofluorescence assays (IFA)were performed at 24 hours post infection (hpi). Infection was confirmed by visualizing presence of viral non-structural protein 4 (nsP4) in the infected cells (Fig. 2A). To detect replication of virus in infected cells, the expression of viral non-structural protein 1 (nsP1) gene was confirmed. Infected cells showed a significant increase in nsP1 gene expression at 24 hpi (Fig. 2B). The viability of the infected cells was confirmed and found to be similar to mock-infected (Fig. S1) Collectively, these data indicate that BMSCs are susceptible to CHIKV infection.

**Fig 1.**
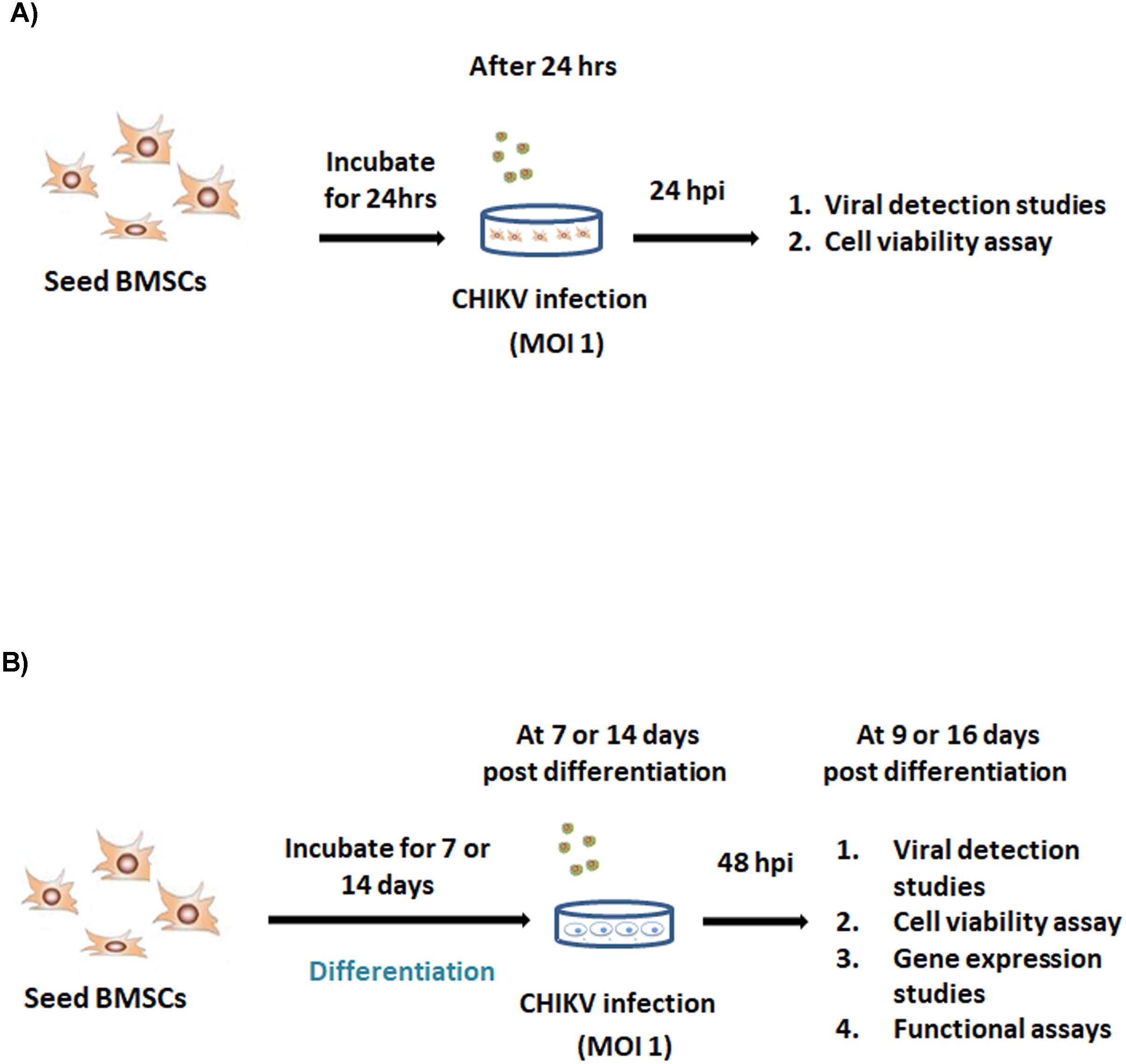
Experimental outline. (A) BMSCs were mock-infected or infected at MOI 1. At 24 hours post infection (hpi), viral detection studies and cell viability assay was performed. (B) BMSCs were differentiated into osteogenic cells. Osteogenic cells at 7 or 14 days post differentiation (dpd) were mock-infected or infected at MOI 1. At 9 or 16 dpd (and at 48 hpi), viral detection studies, cell viability assay, gene expression studies and functional studies were performed.

**Fig 2.**
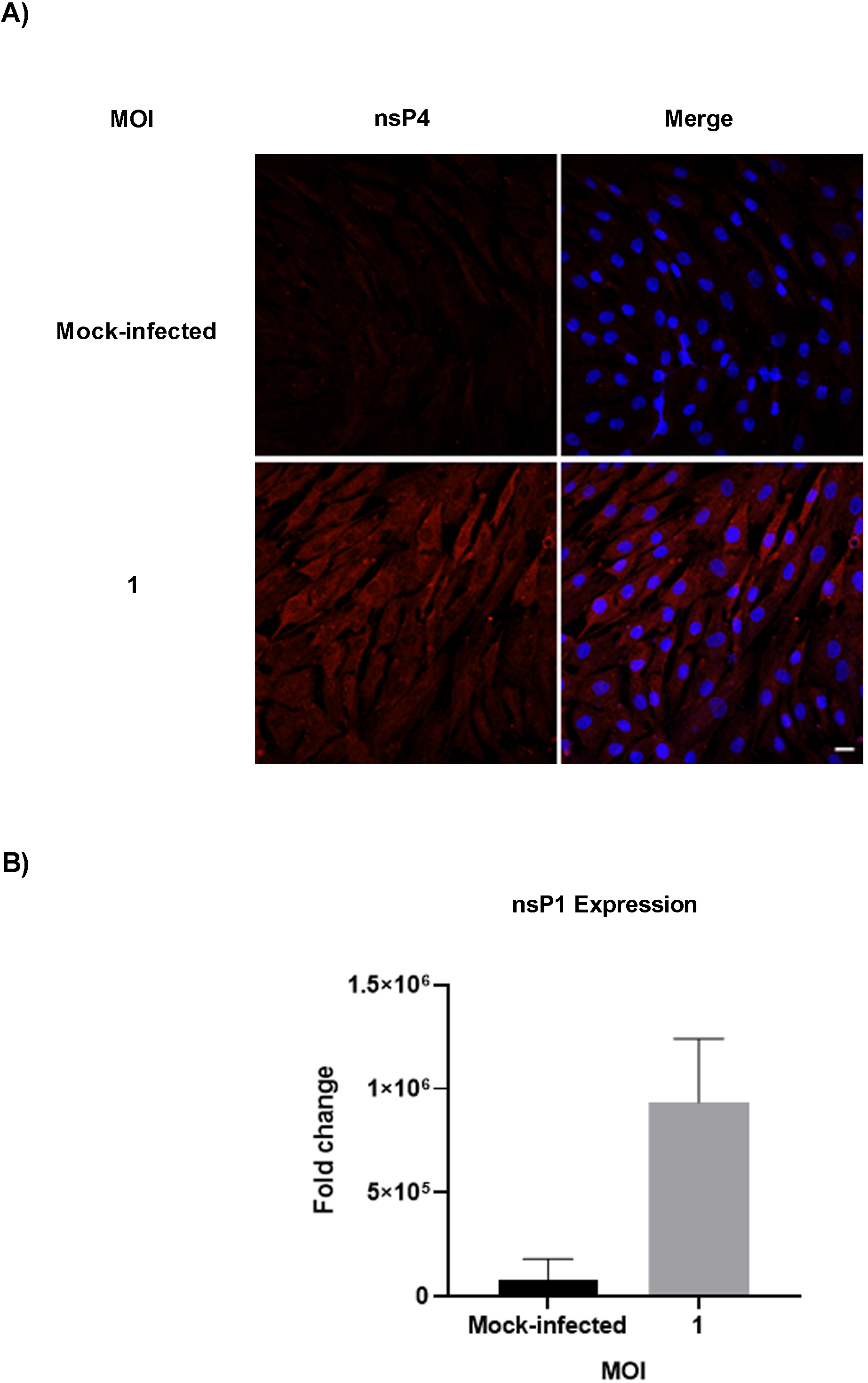
Susceptibility of BMSCs to CHIKV infection. BMSCs were mock-infected or infected with CHIKV at MOI 1 (A) Representative immunofluorescence images of cells fixed and immuno-stained at 24 hpi with antibodies to the viral nsP4 protein (red) and Hoechst nuclear stain (blue). (B) Viral (nsP1) gene expression was quantified by qRT-PCR at 24 hpi. Graph shows fold change in gene expression. Fold change was calculated by 2^-ΔΔCt method. Peptidylprolyl isomerase A (PPIA) was used as a housekeeping gene. (n=3). Error bars show mean ±SEM. Magnification=200x total magnification. Scale bar = 20μm.

### CHIKV infection interferes with gene expression and function of osteogenic cells BMSC-derived osteogenic cells are permissive to CHIKV infection

As mentioned earlier, osteogenic differentiation of BMSCs plays an important role in bone homeostasis. In addition, viral infection has been associated with altered function of these cells [22]. Therefore, we next investigated the impact of CHIKV infection on these cells. BMSCs were differentiated to osteogenic cells and then infected at 7 or 14 days post differentiation (dpd) as outlined in Fig. 1B. Osteogenic differentiation was confirmed by ALP production and matrix mineralization, critical features osteogenic cell function [13, 16]. Production of ALP and the presence of red stained calcium nodes in differentiated cells at 9 and 16 dpd respectively confirmed osteogenic differentiation of BMSCs (Fig 3A and B). To detect the presence of infection in osteogenic cells, we performed IFA at 48 hpi. Infection was confirmed by visualizing presence of viral non-structural protein 4 (nsP4) in the infected cells (Fig. 4A and B). Viral replication in infected cells was confirmed by gene expression of nsP1. A significant increase in nsP1 expression was observed in infected cells at 48 hpi (Fig. 4C and D). The viability of the osteogenic cells was confirmed and found to be similar to that of mock-infected cells at 48 hpi (Fig. S2). Collectively, these results demonstrate that BMSC-derived osteogenic cells are susceptible to CHIKV infection.

**Fig 3.**
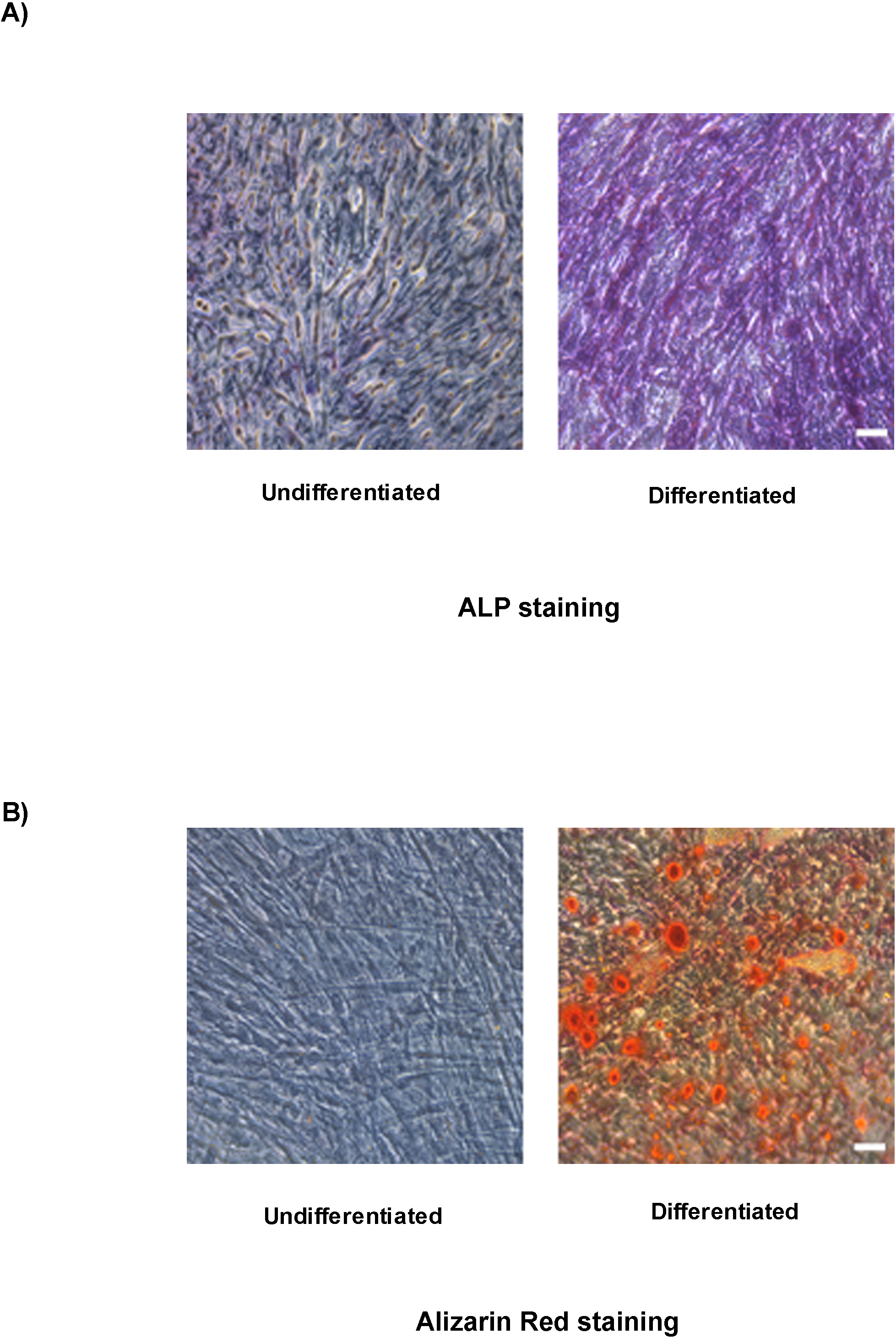
Osteogenic differentiation of BMSCs to osteogenic cells. Representative brightfield images of differentiated and undifferentiated BMSCs, stained with (A) ALP staining and (B) Alizarin Red staining at 9 and 16 dpd. Magnification=200× total magnification. Scale bar = 0.2μm.

**Fig 4.**
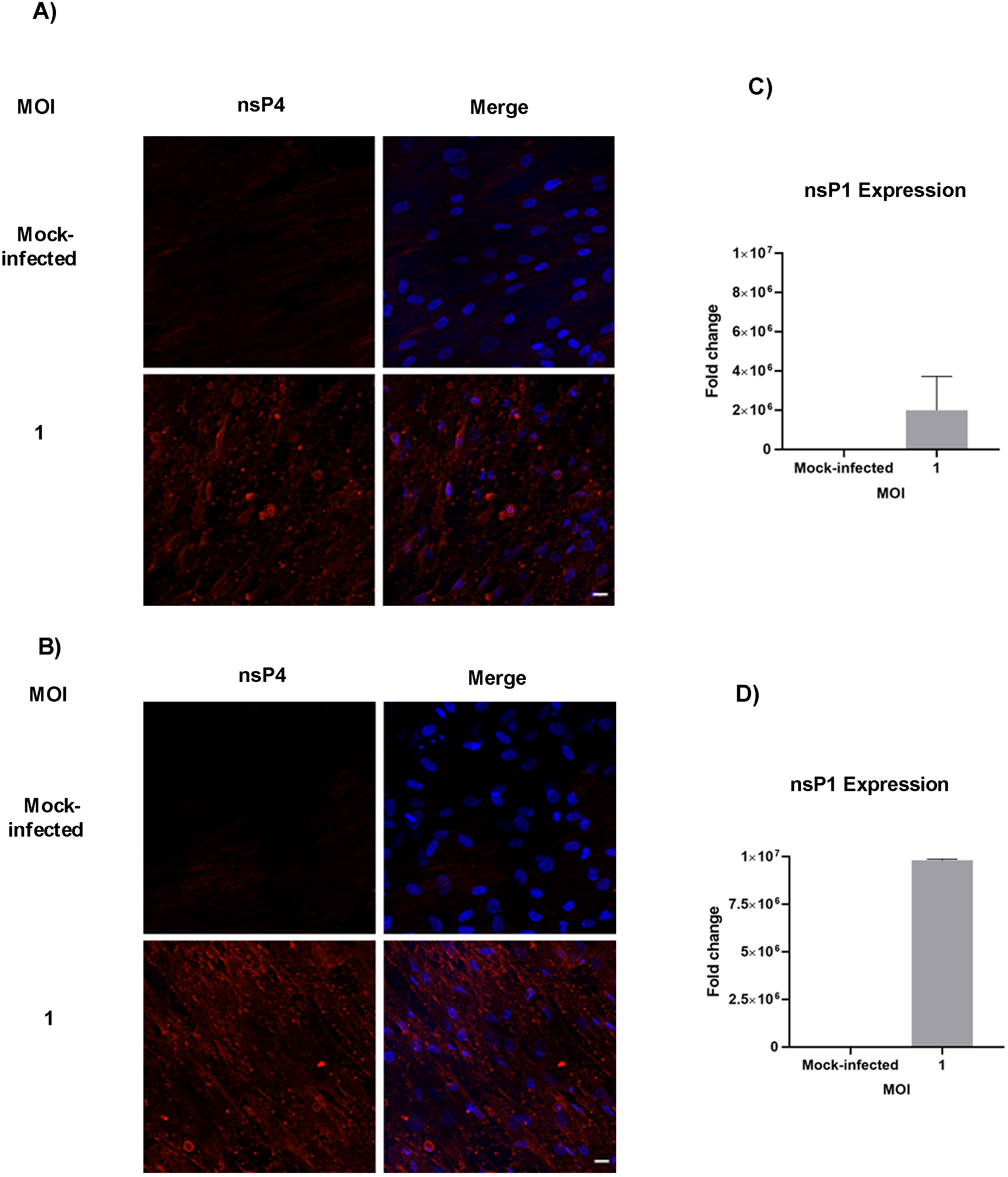
Susceptibility of osteogenic cells to CHIKV infection. Osteogenic cells at 7 or 14 dpd were mock-infected or infected with CHIKV at MOI 1. (A) Representative immunofluorescence images of cells fixed and immuno-stained at 9 or (B) 16 dpd (and at 48 hpi) with antibodies to the viral nsP4 protein (red) and Hoechst nuclear stain (blue). (C) Viral nsP1 gene expression was quantified by qRT-PCR at 9 dpd or (D) 16 dpd (and at 48 hpi). Bar graph shows fold change in gene expression in infected cells compared to mock-infected control. Fold change was calculated by 2^-ΔΔCt method. HPRT1 was used as a housekeeping gene. n=3. Error bars show ±SEM. (D) Representative bright field images of morphological analysis of BMSCs at 7 dpi. n=3. Magnification=200× total magnification. Scale bar = 20μm.

### CHIKV infection decreases gene expression of RUNX2 in osteogenic cells

During differentiation, there is increased expression of osteogenic marker genes including collagen type 1 alpha 1 (COL1A1), ALP, and the marker of early osteogenesis, RUNX2 [5, 15]. Recent studies show that viral infection of osteogenic cells can alter gene expression of osteogenic markers [22]. To determine the effect of CHIKV infection on the gene expression of these markers, osteogenic cells were infected (Fig 1B) and qRT-PCR was performed at 48 hpi (at 9 and 16 dpd) (Fig. 5). Interestingly, a significant decrease in gene expression of RUNX2 was observed in CHIKV infected cells at 16 dpd (Fig. 5B). However, no significant change was observed in the expression of other marker genes in the presence of CHIKV infection (Fig. 5 A, C-F). Thus, these results revealed that CHIKV infection impairs gene expression of the major osteogenic regulator, RUNX2.

**Fig 5.**
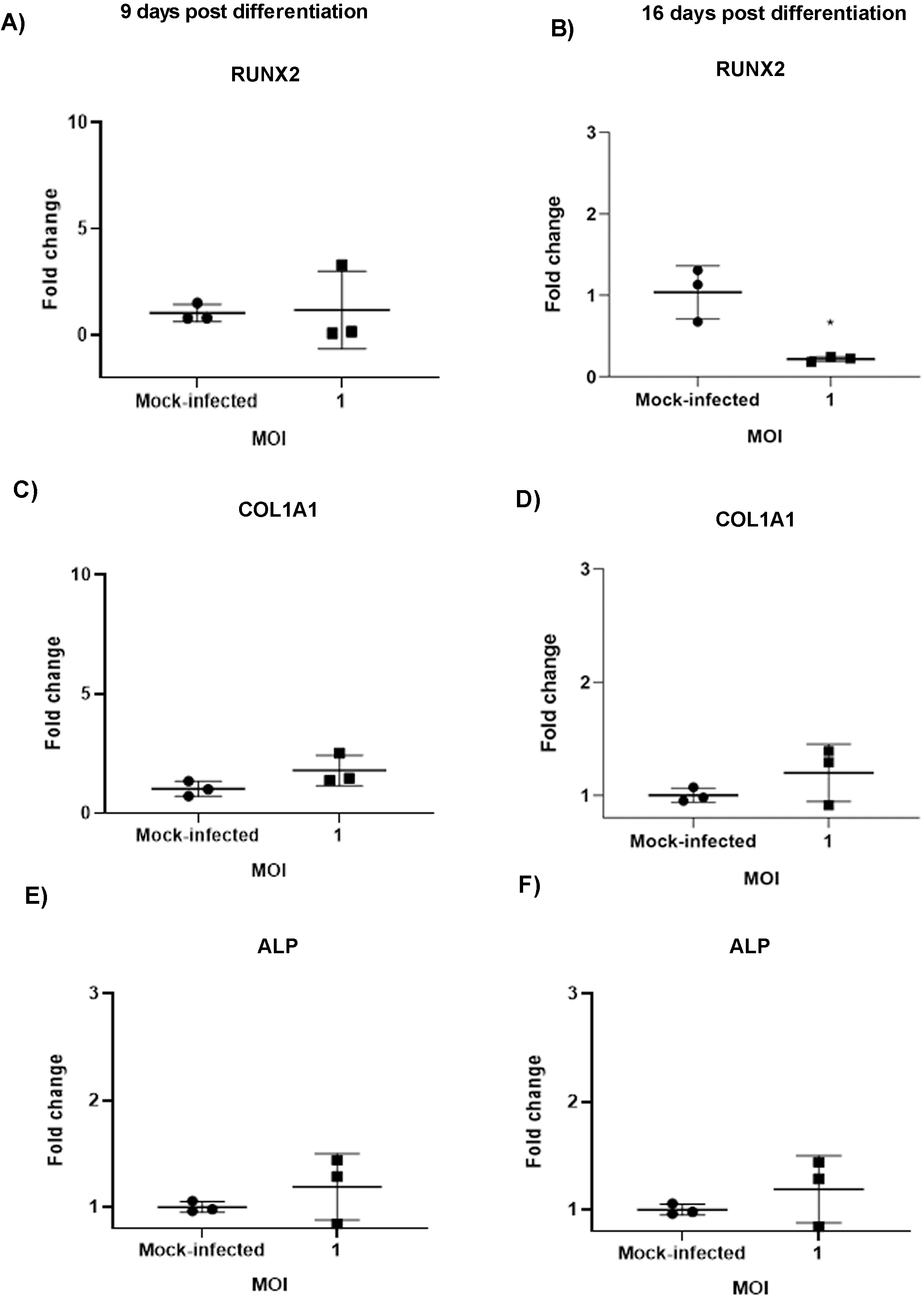
Effect of CHIKV infection on gene expression of osteogenic marker genes. Osteogenic cells at 7 or 14 dpd were mock-infected or infected with CHIKV at MOI 1. (A,C,E) Gene expression of the osteogenic marker genes (RUNX2, COL1A1, ALP) was determined by qRT-PCR at 9 or (B,D,F) 16 dpd (and at 48 hpi). Graph shows fold change in gene expression. Fold change was calculated by 2^-ΔΔCt method. Peptidylprolyl isomerase A (PPIA) was used as a housekeeping gene. (n=3). Error bars show mean ±SEM. Significant changes are represented as p-values (*p<0.05)

### CHIKV infection impairs function of osteogenic cells

As earlier mentioned, ALP is produced during osteogenic differentiation [13]. The production and activity of the protein can be evaluated by well-established assays [13, 16]. A recent study with Zika virus (ZIKV) showed that viral infection of osteogenic cells impaired ALP activity during differentiation [22]. Similarly, we sought to investigate the effect of CHIKV infection on ALP production and activity. Differentiated osteogenic cells were infected for 48 h (at 9 or 16 dpd) and ALP staining and ALP activity assays were performed. A significant reduction in the ALP staining and activity was observed in infected cells at 9 dpd (Fig. 6A and B). However, no significant change in ALP staining or activity was observed in infected cells at 16 dpd (data not shown).

**Fig 6.**
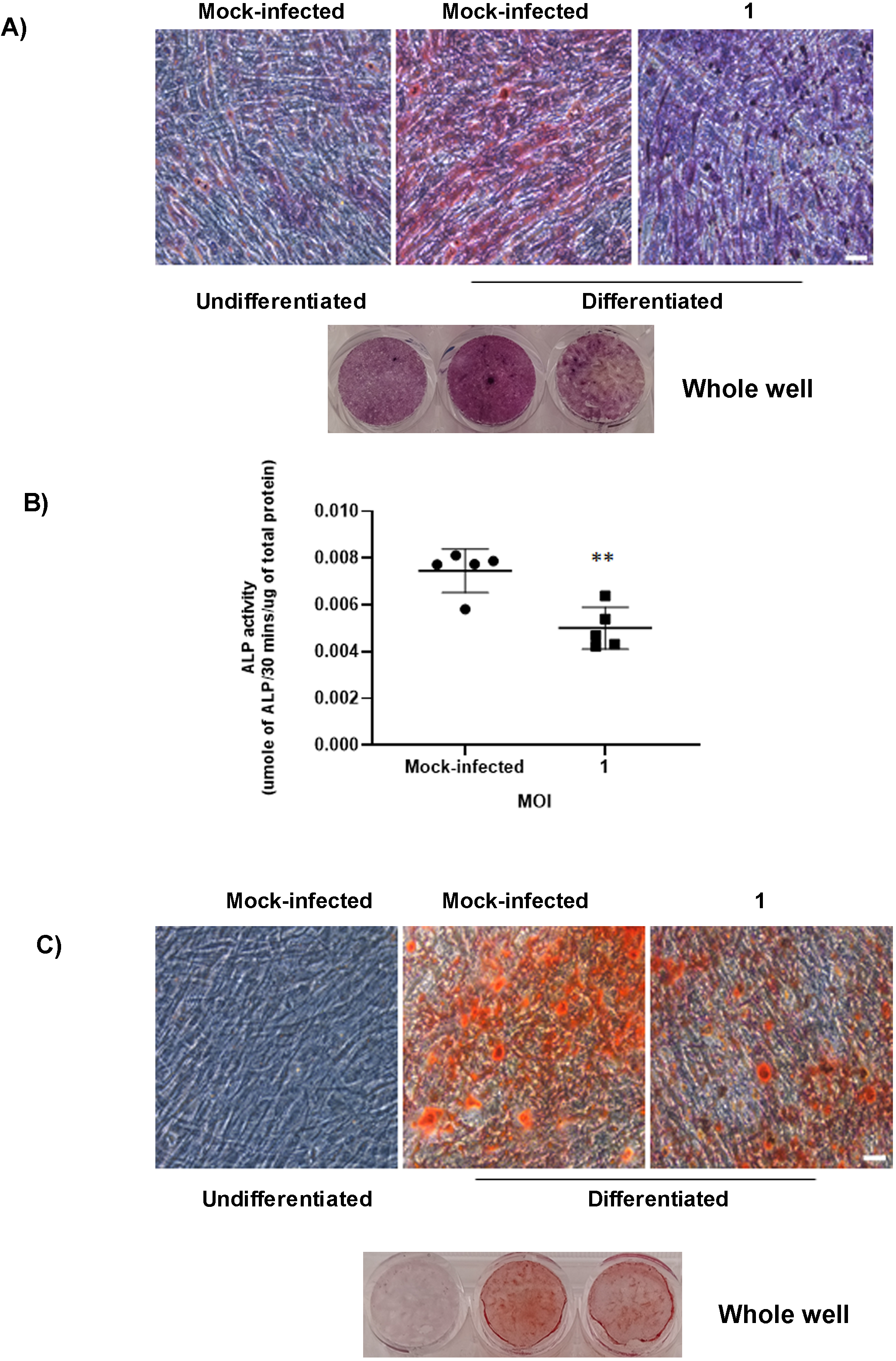
Effect of CHIKV infection on the function of osteogenic cells. Osteogenic cells at 7 or 14 dpd were mock-infected or infected with CHIKV at MOI 1. (A) Representative bright field images of mock-infected or infected cells stained by ALP stain to detect ALP production by the osteogenic cells at 9 dpd (and at 48 hpi). (B) Bar graphs show ALP activity detected by ALP activity assay at 9 dpd (and at 48 hpi). ALP levels were normalized against total protein. (C) Representative brightfield images of mock-infected or infected osteogenic cells, stained with Alizarin Red at 16 dpd (and at 48 hpi). n=5. Error bars show ±SEM. Significant changes are represented as p-values (*p<0.05) Magnification=100X total magnification. Scale bar=0.2 μm.

Osteogenic cells produce calcium phosphate crystals for matrix mineralization during osteogenic differentiation [13]. These crystals can be detected by Alizarin Red staining assay [13]. To determine the effect of CHIKV infection on matrix mineralization, osteogenic cells were infected as outlined in Fig 1B and the staining assay was performed at 9 and 16 dpd (at 48 hpi). A significant decrease in production of calcium phosphate crystals was observed in infected cells at 16 dpd (Fig.6C). However, no significant change in production of calcium phosphate crystals was observed in infected cells at 9 dpd (data not shown). This difference is likely due to calcium phosphate deposition being a later event during osteogenesis [13]. In support of that we did not detect much difference in calcium phosphate deposits between mock-infected undifferentiated and differentiated cells at 9 dpd. Together, these results demonstrate for the first time that CHIKV infection impairs the function of BMSC-derived osteogenic cells.

## Discussion

The involvement of bone pathology has been associated with CHIKV infection, however, the mechanism of bone pathology is unclear [12, 19]. BMSCs are non-hematopoietic multipotent stem cells which can differentiate into osteogenic cells thereby playing an essential role in bone homeostasis [6, 26]. In the current study we show that CHIKV can infect BMSCs and BMSC-derived osteogenic cells and affect function thereby providing a potential mechanism of viral induced bone pathology.

A few reports detected that BMSCs are susceptible to viral infection and this can affect the function and properties of these cells. In a study by Meisel et al, it was reported that BMSCs are susceptible to cytomegalovirus (CMV) infection leading to inhibition of the immunosuppressive properties of BMSCs [20]. However, susceptibility of these cells to arthritic viral infections has not been reported. To our knowledge, this is the first demonstration that these cells are susceptible to CHIKV infection. In this study, we show that BMSCs support viral replication and protein expression. The viability of the infected cells was confirmed to be similar to the mock-infected at the time of study. Interestingly, these cells are susceptible even at very low MOI (data not shown). Studies show that BMSCs migrate to the joint space during injury or disease, so it would be interesting to investigate whether CHIKV can infect BMSCs in vivo and whether infection can affect in vivo osteogenesis [24].

One of the important functions of BMSCs is their ability to differentiate into osteogenic cells and contribute in bone homeostasis [14]. The major hallmarks of osteogenesis are expression of osteogenic marker genes, production of ALP and deposition of calcium phosphate crystals [13, 15]. The most studied markers are COL 1A1, which forms the organic part of the bone, RUNX2, the major regulator of osteogenesis and ALP, essential in matrix mineralization [5, 15]. In our study, we have differentiated BMSCs into osteogenic cells. Osteogenesis was confirmed by ALP production at 7 dpd and calcium phosphate deposition at 14 dpd. This is likely as production of ALP starts early during osteogenesis whereas, the calcium phosphate deposition occurs at a later stage during the process [13]. Prior studies have shown that virus infection can impair function of osteogenic cells [22]. So, we studied the effect of CHIKV infection on the function of these cells. We infected the cells at 7 or 14 dpd and performed the viral detection studies, gene expression studies and functional assays at 48 hpi. We have selected these time points based on literature study [15, 22]. We report that BMSCs-derived osteogenic cells are susceptible to CHIKV infection. We also confirmed the viability of the infected cells as similar to mock-infected. As observed previously, osteoblasts are susceptible to CHIKV, thus, our results are consistent with the prior report [23]. We also demonstrate that infection inhibited gene expression of RUNX2. As been reported previously with a flavivirus, ZIKV infection induced a similar reduction in the gene expression of RUNX2 in infected cells [22]. Thus, further studies are necessary to unravel the mechanism of this infection induced decrease in gene expression. However, Mumtaz et al in their study also reported a decreased ALP gene expression at 11 dpi, whereas, our study showed no significant change in expression of ALP in infected cells. Similarly, no change in expression of COL1A1 was observed in our study. A likely reason for this data may be due to selection of 7 and 14 dpd for our study. It is possible that infection can cause changes in ALP or COL1A1 expression at other stages during differentiation.

During osteogenesis, ALP produced by the osteogenic cells dephosphorylates inorganic pyrophosphate producing inorganic phosphate that binds with calcium to form hydroxyapatite during matrix mineralization [10]. Studies show that an imbalance in ALP production can lead to bone pathology [2]. In our study, we observed that CHIKV infection in osteogenic cells perturbs the production and activity of ALP at 9 dpd. A similar study by Mumtaz et al confirms our result where they reported a decrease in ALP activity in presence of ZIKV infection [22]. Calcium phosphate deposition is essential for matrix mineralization which provides strength and support to the bone [10]. An imbalance in calcium phosphate deposition can also result in diseased condition [3]. To our knowledge, this is the first study that shows CHIKV infection impairs calcium phosphate deposition in infected cells at 16 dpd. However, no change was observed at 9 dpd which is likely as we did not observe any change between mock-infected, undifferentiated and differentiated cells at this time point. Thus, our in vitro model allowed us to conclude that infection lead to the impaired function of osteogenic cells as observed by the reduction in ALP production and activity and matrix mineralization. As bone pathology is associated with CHIKV infection, hence, this in part can explain the arthritic like conditions and joint pain observed in CHIKV infection [12, 19].

However, matrix mineralization is also associated with production of extracellular matrix (ECM) proteins like fibronectin, vitronectin, laminin, osteopontin, osteonectin [9]. Thus, studying the expression of these proteins and their interaction with viral proteins during infection would provide a better understanding of viral pathogenesis in this context.

Additionally, in vivo differentiation of BMSCs into osteogenic cell lineages depends on interaction of these cells with other cells in the joint tissue [13], but our current in vitro model do not mimic the joint microenvironment. Hence, co-culturing of BMSCs with other CHIKV permissive cells from joint tissue would lead to a better understanding of the pathogenesis of CHIKV infection. Additionally, the receptor activator of nuclear factor kappa-B ligand (RANKL)/osteoprotegerin (OPG) ratio is a key determinant of bone mass and strength, and thus, plays an important role in bone homeostasis [4]. Thus, future studies can be targeted to study protein and gene expression of RANKL and OPG in the presence of CHIKV infection.

Based on our observations, we propose BMSCs as a suitable model for studying bone homeostasis in the context of CHIKV infection. Most importantly, we confirm that infection produces functionally altered osteogenic cells and likely lead to dysregulation in the dynamics of bone homeostasis often observed during in this disease.

## Materials and Methods

### Cell culture, Compounds and Virus

Human BMSCs were purchased from Roosterbio (USA). CHIKV (181/25 clone) was purchased from BEI Resources.

BMSCs were infected at a MOI 1 and incubated for 1 h at 37°C, 5% CO_2_ (Fig. 1A). At one-hour post infection (hpi), cells were washed with DPBS (Corning, USA), replenished with Rooster Basal MSC (RBM) media (RoosterBio, USA) and incubated at 37 °C, 5% CO_2_ for 24 h.

For differentiation study, BMSCs were treated with osteogenic media containing mesenchymal stem cell expansion medium (Millipore, USA) and supplemented with 10 mM beta glycerophosphate (Sigma, USA), 0.1 μM dexamethasone (Sigma, USA) and 200 μM ascorbic acid (Sigma, USA) to differentiate into osteogenic cells. The osteogenic cells were infected at 7 or 14 dpd, at a MOI 1 and incubated for 1 h at 37°C, 5% CO_2_ (Fig. 1B). After 1 hpi, cells were washed with DPBS, replenished with osteogenic differentiation media and incubated for 48 h at 37°C, 5% CO_2_. For these experiments, mock-infected undifferentiated cells were negative controls, while mock-infected differentiated cells were positive controls.

### Immunofluorescence assay

BMSCs at a density of 2.5×10^4^ per well were seeded in 4-well plate containing glass cover slips in each well and treated as outlined in Fig. 1A and B. Mock or virus-infected cells were fixed in 4% paraformaldehyde (PFA) (Electron Microscopy Sciences, USA) for 30 min and permeabilized with 0.1% Triton X (Fisher Bioreagents, USA) for 10 mins. Blocking was done in 3% Bovine Serum Albumin-Phosphate Buffered Saline for 1 h. Virus was stained with antibody against CHIKV non-structural protein (nsP4) (kindly provided by Dr. Andres Merits). Following overnight incubation with primary antibody at 4 °C, the cells were stained with Alexa-Fluor 568 goat anti-rabbit IgG (Life Technologies, USA) for 1 h at room temperature. Nuclear staining was done with Hoechst 33642 (Invitrogen, USA) for 30 min. Images were taken and processed using Zeiss LSM 800 with Airyscan.

### Cell viability assay by Live and Dead staining

BMSCs at a density of 3×10^3^ per well were seeded in a 96-well plate and treated as outlined in Fig. 1A and B. BMSCs or osteogenic cells were stained with calcein acetoxymethylester (calcein AM) (Biotium, California, USA) and ethidiumhomodimer-2 (Biotium, California, USA), at a concentration of 2 μM and 4 μM respectively and incubated for 30 min at 37°C, 5% CO_2_. Calcein AM stain viable cells with bright green fluorescence [17]. Ethidium homodimer stain nonviable cells with red fluorescence but cannot penetrate viable cells [17]. After 30 min incubation, samples were imaged using a High Content Analysis System, Operetta CLS^™^ and the percentage of viable cells were calculated using the Harmony^®^ analysis software. Staurosporine was taken as control.

### ALP staining and ALP activity assay

ALP production by the osteogenic cells was examined by ALP staining assay [1]. BMSCs at a density of 3×10^4^ per well were seeded in a 24-well plate and treated as outlined in Fig. 1B. At 9 and 16 dpd (and at 48 hpi), ALP staining was performed with the osteogenic cells using ALP leukocyte kit (Sigma-Aldrich, USA) according to manufacturer’s instructions. Images were taken using a DM1/MC120 microscope (Leica, Germany) at 20× magnification.

ALP enzyme activity was determined by ALP activity assay [16]. BMSCs at a density of 2.5×10^4^ per well were seeded in a 24-well plate and treated as outlined in Fig 1B. At 9 and 16 dpd (and at 48 hpi) the infected osteogenic cells were lysed by freeze-thaw method in a buffer containing Triton X-100 (0.1% v/v) (Fischer Scientific, USA), 1 mM MgCl_2_ (Alfa Aesar, USA), 20 mM Tris (Fischer Scientific, USA). The cell lysate was used to perform ALP activity assays using the ALP activity kit (Sigma-Aldrich, USA) according to the manufacturer’s instructions. The ALP activity assay uses p-nitrophenyl phosphate (pNPP) as substrate which is dephosphorylated by the ALP enzyme forming *p*-nitrophenol which was measured spectrophotometrically at 405 nm in Synergy H1 Hybrid Reader (BioTek) [16]. The total protein content in the infected cells was determined using a BCA protein assay kit (Thermo Scientific, USA) with bovine serum albumin as standard. The ALP activity was expressed as micromole of p-nitrophenol formed per 30 minutes per microgram of total protein (μmol per 30 min per μg protein).

### Alizarin Red staining

BMSCs at a density of 5×10^4^ per well were seeded in a 24-well plate and treated as outlined in Fig. 1B. At 9 and 16 dpd, Alizarin Red staining assay was performed with the osteogenic cells using Alizarin Red solution (EMD Millipore, Germany) according to manufacturer’s instructions. Alizarin Red stains the calcium phosphate deposits formed by matrix mineralization during osteogenic differentiation [13]. All images were taken using DM1/MC120 microscope (Leica, Germany) at 20× magnification.

### Quantitative RT-PCR (qRT-PCR)

BMSCs at a density of 2×10^5^ per well were seeded in a 6-well plate and treated as outlined in Fig. 1A and B. BMSCs or osteogenic cells were collected at time points outlined in Fig. 1A and B and lysed in RLT buffer (Qiagen, Germany) for RNA isolation. RNA isolation was performed using RNeasy Mini kit (Qiagen, Germany) according to the manufacturer’s instructions. RNA was quantified and total RNA was reverse transcribed using a qScript cDNA synthesis kit (Quantabio, USA) according to the manufacturer’s instructions. SYBR Green Real-Time PCR was performed on a StepOnePlus Real-Time PCR System (Thermo Scientific) using SSO Advanced SYBR Green Supermix (Bio-Rad, USA) using the primers for osteogenic marker genes i.e. RUNX2 (5’-TACCTGAGCCAGATGACG-3’ and 5’-AAGGCCAGAGGCAGAAGT-3’), COL1A1 (5’-CGAAGACATCCCACCAATC-3’ and 5’-ATCACGTCATCGCACAACA-3’), ALP (5’-TCACTCTCCGAGATGGTGGT-3’ and 5’-GTGCCCGTGGTCAATTCT-3’) (IDT, Coralville, USA) and viral non-structural (nsP1) gene (Applied Biosystems, USA) (5’-GGGCTATTCTCTAAACCGTTGGT-3’ and 5’-CTCCCGGCCTATTATCCCAAT-3’) according to the manufacturer’s instructions with the following conditions: (i) PCR initial activation step, 95 °C for 3 min, 1 cycle, and (ii) three-step cycling, 95 °C for 15s, followed by 60 °C for 1min and 95 °C for 15s, 40 cycles. The fold change in mRNA expression relative to mock-infected samples was calculated using the 2^-ΔΔCt method. Transcript levels were normalized using peptidylprolyl isomerase A (IDT, Coralville, USA).

### Statistical analysis

All statistical analyses were performed using GraphPad Prism 8. Data were represented as the mean ± standard error of the mean (SEM). Significant differences between the experimental groups were determined using Student’s t test. P values < 0.05 are shown.

## Acknowledgments

We thank Dr. Leah Cook for critical reading of the manuscript and valuable suggestions. We thank Janice A. Taylor and James R. Talaska of the Advanced Microscopy Core Facility at the University of Nebraska Medical Center for providing assistance with confocal microscopy. We also thank Dr. Andres Merits for kindly providing the nsP4 antibody. This study was supported by NIH/NIAID 1R21AI140026-01 (SPR & BD) and start-up funds (SPR).

---

Fig S1. Cell viability assay. BMSCs were mock-infected or infected with CHIKV at MOI 1. Percentage of dead cells determined by Live/Dead staining assay at 24 hpi (left panel). Representative microscopic images of BMSCs at 24 hpi, stained with calcein AM (stains the viable cells) and ethidium homodimer (stains the non-viable cells) (right panel). Magnification=200× total magnification. Scale bar = 100μm.

Fig S2. Cell viability assay. Osteogenic cells at 7 or 14 dpd were mock-infected or infected with CHIKV at MOI 1. (A) Percentage of dead cells determined by Live/Dead staining assay at 9 dpd (left panel). Representative microscopic images of osteogenic cells at 9 dpd (and at 48 hpi), stained with calcein AM (stains the viable cells) and ethidium homodimer (stains the non-viable cells) (right panel). (B) Percentage of dead cells determined by Live/Dead staining assay at 16 dpd (left panel). Representative microscopic images of osteogenic cells at 16 dpd (and at 48 hpi) (right panel). Magnification=200× total magnification. Scale bar = 100μm

